# Structural rearrangements drive extensive genome divergence between symbiotic and free-living *Symbiodinium*

**DOI:** 10.1101/783902

**Authors:** Raúl A. González-Pech, Timothy G. Stephens, Yibi Chen, Amin R. Mohamed, Yuanyuan Cheng, David W. Burt, Debashish Bhattacharya, Mark A. Ragan, Cheong Xin Chan

## Abstract

Symbiodiniaceae are predominantly symbiotic dinoflagellates critical to corals and other reef organisms. *Symbiodinium* is a basal symbiodiniacean lineage and includes symbiotic and free-living taxa. However, the molecular mechanisms underpinning these distinct lifestyles remain little known. Here, we present high-quality *de novo* genome assemblies for the symbiotic *Symbiodinium tridacnidorum* CCMP2592 (genome size 1.3 Gbp) and the free-living *Symbiodinium natans* CCMP2548 (genome size 0.74 Gbp). These genomes display extensive sequence divergence, sharing only ~1.5% conserved regions (≥90% identity). We predicted 45,474 and 35,270 genes for *S. tridacnidorum* and *S. natans*, respectively; of the 58,541 homologous gene families, 28.5% are common to both genomes. We recovered a greater extent of gene duplication and higher abundance of repeats, transposable elements and pseudogenes in the genome of *S. tridacnidorum* than in that of *S. natans*. These findings demonstrate that genome structural rearrangements are pertinent to distinct lifestyles in *Symbiodinium*, and may contribute to the vast genetic diversity within the genus, and more broadly in Symbiodiniaceae. Moreover, the results from our whole-genome comparisons against a free-living outgroup support the notion that the symbiotic lifestyle is a derived trait in, and that the free-living lifestyle is ancestral to, *Symbiodinium*.

## Introduction

Symbiodiniaceae are dinoflagellates (Order Suessiales) crucial for coral reefs because of their symbiotic relationship with corals and diverse marine organisms. Although these dinoflagellates do not display evident morphological diversity, their extensive genetic variation is well-recognised, prompting the recent systematic revision to family status^1,2^. Sexual reproduction stages have not been directly observed in Symbiodiniaceae, but the presence of a complete meiotic gene repertoire suggests that they are able to reproduce sexually^3–5^. The potential sexual reproduction of Symbiodiniaceae has been used to explain their extensive genetic variation^6–10^.

The genetic diversity in Symbiodiniaceae is in line with their broad range of symbiotic associations with other organisms, covering a broad spectrum depending on host specificity, transmission mode and permanence in the host^11,12^. Furthermore, some taxa are considered free-living because they have been found only in environmental samples, and in experiments fail to infect potential hosts^13–15^.

The basal lineage of Symbiodiniaceae (formerly clade A) consists of two monophyletic groups, one of which has been revised as *Symbiodinium sensu stricto*^2,16^. *Symbiodinium* (as revised) includes a wide range of mutualistic, opportunistic and free-living forms. *Symbiodinium tridacnidorum*, for instance, encompasses isolates in *ITS2*-type A3 that are predominantly symbionts of giant clams in the Indo-Pacific Ocean^2^. Although the nature of this symbiosis is extracellular, they can also establish intracellular symbiosis with cnidarian hosts both in experimental settings and in nature^17^. On the other hand, *Symbiodinium natans* (the type species of the genus) is free-living. *S. natans* occurs frequently in environmental samples, exhibits a widespread distribution and, thus far, has not been shown to colonise cnidarian hosts^2,18^.

Symbiosis, or the lack thereof, has been predicted to impact genome evolution of Symbiodiniaceae^12^. Most symbiotic Symbiodiniaceae are thought to be facultative to some extent, with the potential to shift between a free-living motile stage (*i.e.* mastigote form) and a spherical symbiotic stage (*i.e.* coccoid form). The genomes of facultative and recent intracellular symbionts and parasites are usually very unstable, with extensive structural rearrangements, intensified activity of transposable elements (TEs) and exacerbated gene duplication that leads to the accumulation of pseudogenes^19,20^. Symbiotic Symbiodiniaceae are thus expected to display similar genomic features.

In this study, we present draft *de novo* genome assemblies of *S. tridacnidorum* CCMP2592 and *S. natans* CCMP2548. Using a comparative genomic approach, we found extensive genome-sequence divergence and few shared families of predicted genes between the two species. A greater extent of gene duplication, and the higher abundance of TEs and pseudogenes in *S. tridacnidorum* relative to *S. natans* suggest that duplication and transposition underpin genome divergence between these species.

## Results

### Genome sequences and predicted genes of *S. tridacnidorum* and *S. natans*

The genome sequences of *S. tridacnidorum* CCMP2592 and *S. natans* CCMP2548 were assembled *de novo* using both short- and long-read sequence data (**Error! Reference source not found.**, Supplementary Table 1). The estimated genome size is 1.29 Gbp for *S. tridacnidorum*, and 0.74 Gbp for *S. natans* (Supplementary Table 2); the latter is the smallest reported for any Symbiodiniaceae genome to date. Using an integrative gene-prediction workflow tailored for dinoflagellate genomes (see Methods), we predicted 45,474 high-quality gene models in *S. tridacnidorum*, and 35,270 in *S. natans* (**Error! Reference source not found.**). The gene repertoire for each genome is more complete (85.15% and 83.41% recovery of core conserved eukaryote genes^21^ in *S. tridacnidorum* and *S. natans*, respectively) than other *Symbiodinium* genomes (<79% recovery; Supplementary Figure 1).

**Fig. 1.**
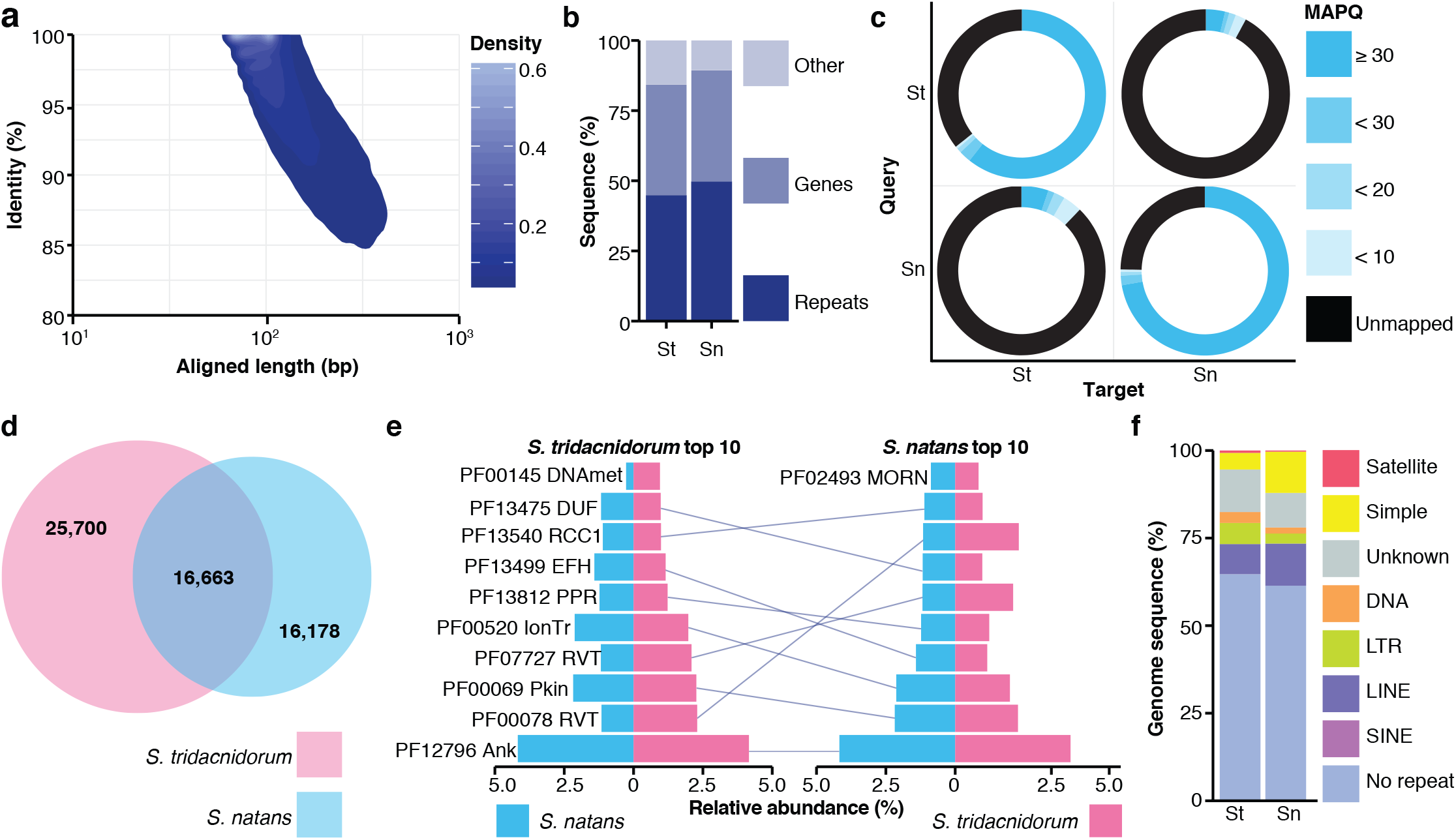
Comparison of *S. tridacnidorum* and *S. natans* genomes. **(a)** Density polygon of the similarity between aligned genome sequences of *S. tridacnidorum* and *S. natans* as a function of the length of the aligned region in the query sequence. **(b)** Proportion of distinct genome features (by sequence length) among the aligned regions between the two genomes. Overlap of the sequences with similarity between both genomes with predicted genes and repetitive elements. **(c)** Mapping rate of filtered read pairs generated for each species against the assembled genomes of itself and of the counterpart. ‘St’: *S. tridacnidorum*, ‘Sn’: *S. natans*. **(d)** Homologous gene families for the two genomes, showing the number of shared families and those that are exclusive to each genome. **(e)** Top ten most-abundant protein domains recovered, sorted in decreasing relative abundance (from bottom to top) among proteins of *S. tridacnidorum* (left) and those of *S. natans* (right). The abundance for each domain in both genomes is shown in each chart for comparison. Domains common among the top ten most abundant for both species are connected with a line between the charts. ‘MORN’: MORN repeat, ‘RCC1’: Regulator of chromosome condensation repeat, ‘RVT’: reverse transcriptase, ‘DUF’: domain of unknown function, ‘PPR’: pentatricopeptide repeat, ‘EFH’: EF-hand, ‘IonTr’: ion transporter, ‘Pkin’: protein kinase, ‘Ank’: ankyrin repeat, ‘DNAmet’: C-5 cytosine-specific DNA methylase. **(f)** Composition of sequence features for each of the two genomes, showing the percentage of sequences (by length) associated with distinct types of repetitive elements. ‘St’: *S. tridacnidorum*, ‘Sn’: *S. natans*.

### Genomes of *S. tridacnidorum* and *S. natans* are highly divergent

The genomes of *S. tridacnidorum* and *S. natans* are highly dissimilar from one another (Fig. 1). Only 14.70 Mbp (1.33%) of the genome sequence of *S. tridacnidorum* aligned to 11.84 Mbp (1.55%) of that of *S. natans* at 90% identity or greater. Most aligned genomic regions are short (<100 bp, Fig. 1a). About half of these regions represent repeats, and another ~40% represent genic regions that are common to both species (Fig. 1b). We observed a low mapping rate (<15%) of read pairs from one genome dataset against the genome assembly of the counterpart, and *vice versa* (Fig. 1c). Using all predicted genes, we inferred 58,541 gene families (including 26,649 single-copy genes), many of which are exclusive to each species (Fig. 1d), *e.g.* 25,700 are specific to *S. tridacnidorum*. However, the predominant gene functions are conserved, as shown by the top ten most abundant protein domains encoded in the genes from both species (Fig. 1e). The composition of repetitive elements differs between the two genomes. Simple repeats and long interspersed nuclear elements (LINEs), for instance, are in smaller proportion in the genome of *S. tridacnidorum* than they are in that of *S. natans* (Fig. 1F). Conversely, long terminal repeats (LTRs) and DNA transposons are more prominent in *S. tridacnidorum*.

### Duplication events and transposable elements contribute to the divergence between *S. tridacnidorum* and *S. natans* genomes

We further assessed the distinct genome features in each species that may have contributed to the discrepancy in genome sizes. Specifically, we assessed, for each feature, the ratio (Δ) of the total length of the implicated sequence regions in *S. tridacnidorum* to the equivalent length in *S. natans* (Fig. 2). The genome size estimate for *S. tridacnidorum* is 1.74 times larger than that for *S. natans* (Supplementary Table 2); we use this ratio as a reference for comparison. Most of the examined genome features span a larger region in the genome of *S. tridacnidorum*, as expected. The Δ for each inspected genic feature (even for exons and introns separately), approximates 1.74. However, six features related to duplicated genes and repetitive elements have Δ > 1.74. This observation suggests that gene duplication and repeats likely expanded in *S. tridacnidorum* (and/or contracted in *S. natans*), contributing to the genome-size discrepancy.

**Fig. 2.**
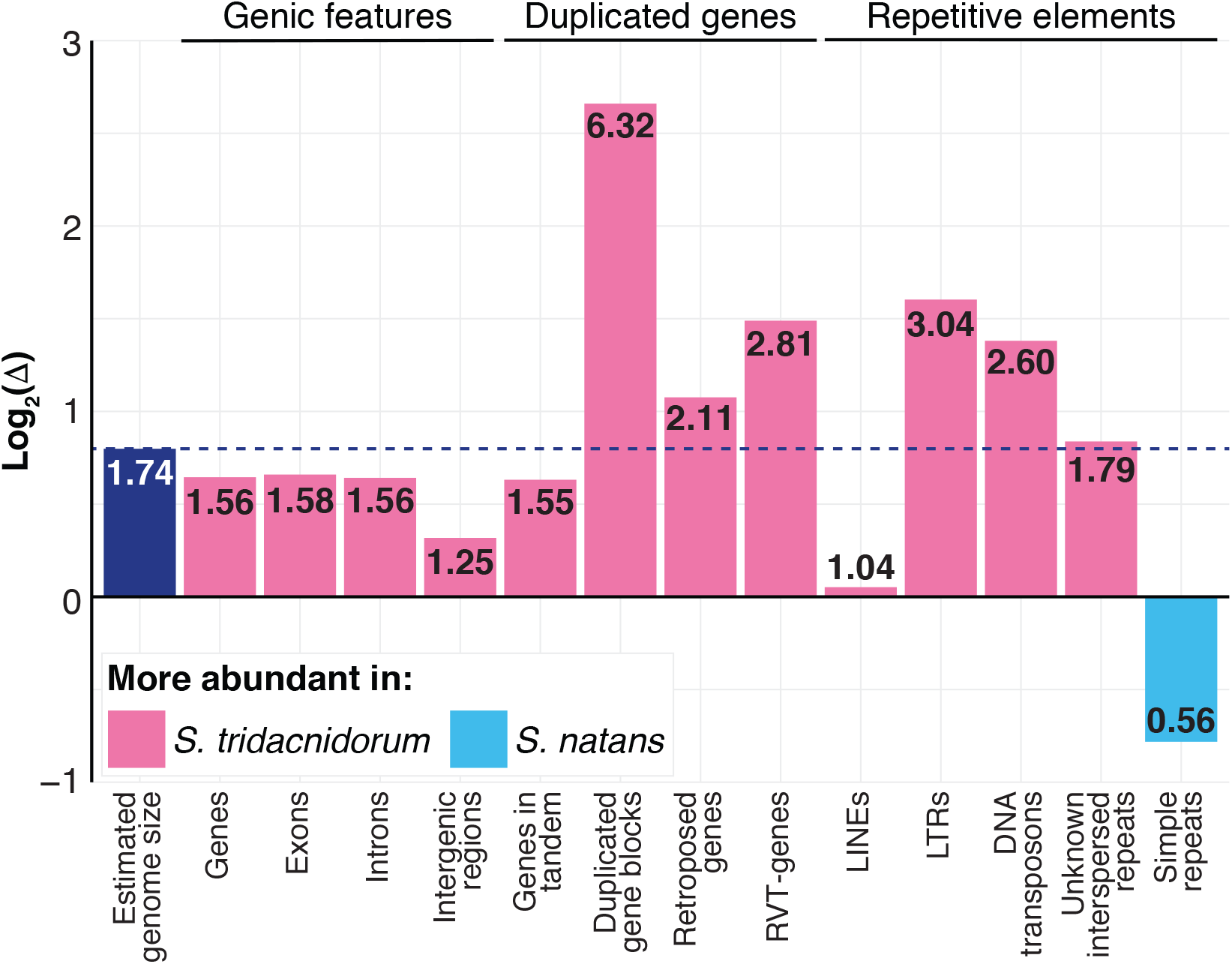
Contribution of genomic features to the distinct composition of *S. tridacnidorum* and *S. natans* genomes. Each genome feature was assessed based on the ratio (Δ) of the total length of the implicated sequence region in *S. tridacnidorum* to the equivalent length in *S. natans*, shown in log_2_-scale. The ratio of the estimated genome sizes is shown as reference (marked with a dashed line). The untransformed Δ for each feature is shown in its corresponding bar. A genome feature with Δ greater than the reference likely contributed to the discrepancy of genome sizes. Bars are coloured based on the genome in which they are more abundant as shown in the legend.

Tandem duplication of exons and genes is common in dinoflagellates, and may serve as an adaptive mechanism to enhance functions relevant for their biology^22,23^. Whereas in some dinoflagellates genes in tandem arrays can have hundreds of copies, *e.g.* up to 5000 copies of the peridinin-chlorophyll a-binding protein (PCP) gene in *Lingulodinium polyedra*^24^, these arrays are not as prominent in the genomes of *S. tridacnidorum* and *S. natans* (Supplementary Figure 2), with the largest array comprising 10 and 13 gene copies, respectively. The 13-gene array in *S. natans* encodes a full-length alpha amylase, whereas the remaining 12 copies are fragments of this gene and likely not functional. On the other hand, the 10-gene block in *S. tridacnidorum* contains genes encoding PCP; of these, seven contain duplets of PCP domains, lending support to the previous finding of the origin of a PCP form by duplication in Symbiodiniaceae^25^; the remaining three copies contain 1, 6 and 14 PCP domains respectively. An additional gene, not part of the tandem array, contains another PCP-duplet. The total 37 individual PCP domains (35 in a gene cluster and two in a separate duplet) supports the earlier size estimation (36 ± 12) of the PCP family in a genome of Symbiodiniaceae^26^. In stark contrast, we only recovered a duplet of PCP domains among all predicted proteins of *S. natans*.

The length of duplicated gene blocks is drastically longer in *S. tridacnidorum* than in *S. natans* (Δ = 6.32; Fig. 2). This observation, and the number of gene-block duplicates in each of the two species, suggests that segmental duplication has occurred more frequently during the course of genome evolution of *S. tridacnidorum*. We found 23 syntenic collinear blocks within the *S. tridacnidorum* genome (*i.e.* within-genome duplicated gene blocks) implicating 242 genes in total. Of these genes, 20 encode protein kinase functions (Supplementary Table 3) that are associated with distinct signalling pathways. In comparison, only five syntenic collinear blocks implicating 62 genes were found in the *S. natans* genome; these genes largely encode functions of cation transmembrane transport, relevant for the maintenance of pH homeostasis. Ankyrin and pentatricopeptide repeats are common in the predicted protein products of duplicated genes in both genomes.

Retroposition is another gene-duplication mechanism known to impact genome evolution of Symbiodiniaceae and other dinoflagellates^22,27^. To survey retroposition in genomes of *S. tridacnidorum* and *S. natans*, we searched for relicts of the dinoflagellate spliced-leader (DinoSL) sequence in upstream regions of all predicted genes. Since the DinoSL is attached to transcribed genes by trans-splicing^28^, genes containing these relicts represent the primary evidence of retroposition into the genome. We found 412 and 252 genes with conserved DinoSL relicts in *S. tridacnidorum* and *S. natans*, respectively. Genes with higher expression levels have been assumed to be more prone to be retroposed into the genome^29^. The identified retroposed genes in the two species encode distinct functions based on the annotated Gene Ontology (GO) terms (Fig. 3a). This observation may be attributed to the preferential expression of functions that are (or were) relevant to each species. For instance, peptide antigen binding (GO:0042605) might be important for host recognition in *S. tridacnidorum*^30^.

**Fig. 3.**
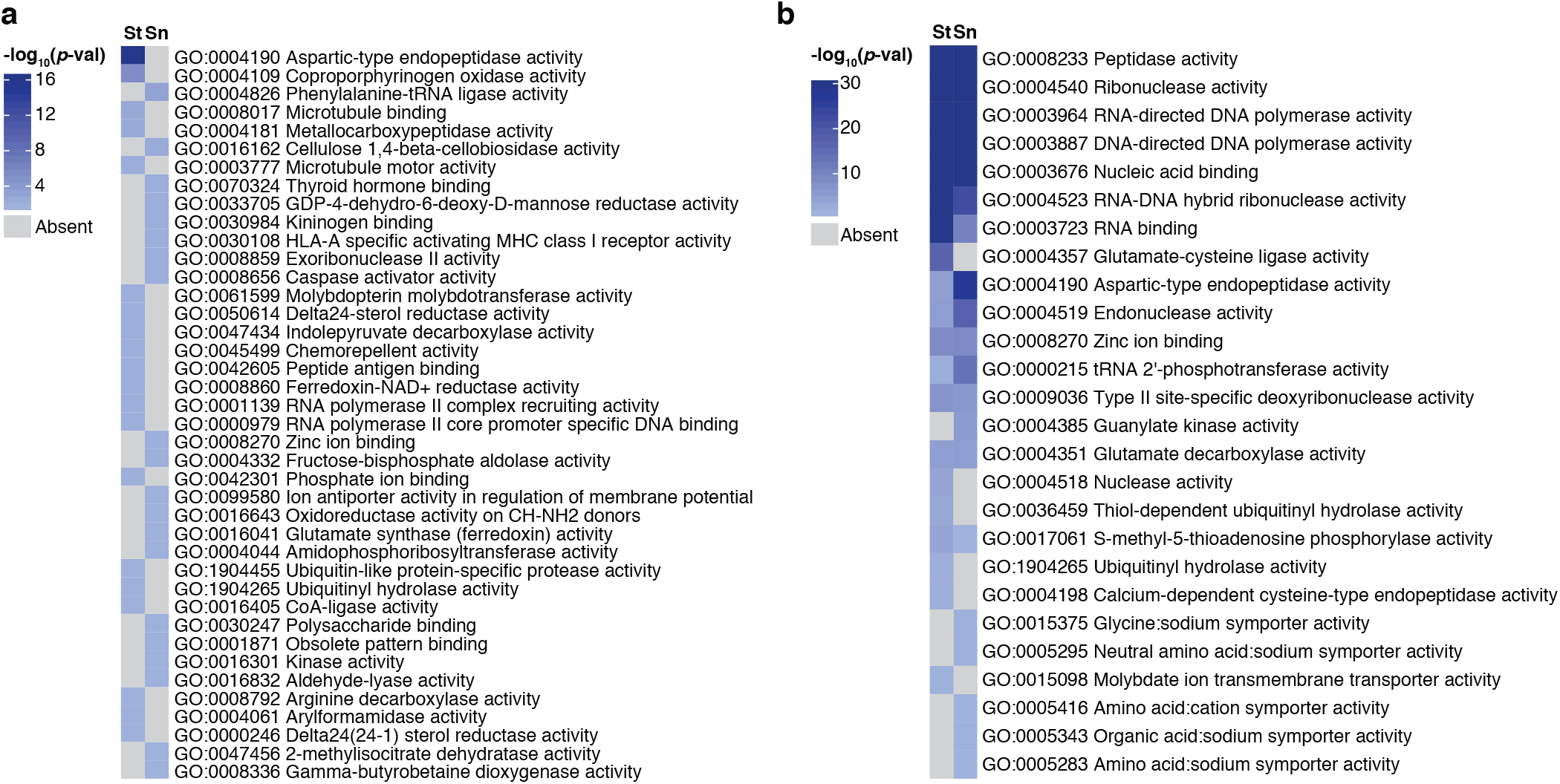
Overrepresented functions in retroposed and RT-genes. GO molecular functions enriched in genes with conserved DinoSL relicts in their upstream regions **(a)** and genes coding for reverse transcriptase domains (RT-genes) **(b)**.

Both retroposition and retrotransposition have been reported to contribute to gene-family expansion in Symbiodiniaceae^31^. Protein domains with functions related to retrotransposition were overrepresented in gene products of *S. tridacnidorum* relative to those of *S. natans* (Supplementary Table 4). However, the reverse transcriptase domains (PF00078 and PF07727) are abundant in both; they were found in 1313 predicted proteins in *S. tridacnidorum* and 591 in *S. natans*.

Retrotransposons can accelerate mutation rate^32^ and alter the architecture of genes in their flanking regions^33^, and may explain the emergence of genes coding for reverse transcriptase domains (RT-genes) in these genomes. Other domains found in these proteins are involved in diverse cellular processes including ubiquitin-mediated proteolysis, DNA methylation, transmembrane transport and photosynthesis (Fig. 3b, Supplementary Table 5). The lack of overlap between functions enriched in genes containing DinoSL relicts and those in RT-genes indicates that retroposition and retrotransposition are independent processes. The abundance of repeats characteristic of TEs (such as LINEs and LTRs; Fig. 2) further supports the enhanced activity of retrotransposition in *S. tridacnidorum*. Although LINEs display high sequence divergence (Kimura distance^34^ 20-30), potentially a remnant from an ancient burst of this type of element common to all Suessiales^3,22^, most LTRs and DNA transposons are largely conserved (Kimura distance < 5), suggesting that they may be active (Fig. 4). We note that these conserved LTRs and DNA transposons were recovered only in our hybrid genome assemblies incorporating both short- and long-read sequence data, and not in our preliminary genome assemblies based solely on short-read data (Supplementary Figure 3, Supplementary Table 6). This indicates that these conserved, repetitive regions can be resolved only using long-read sequence data (Supplementary Figure 4), highlighting the importance of long-read data in generating and assembling dinoflagellate genomes.

**Fig. 4.**
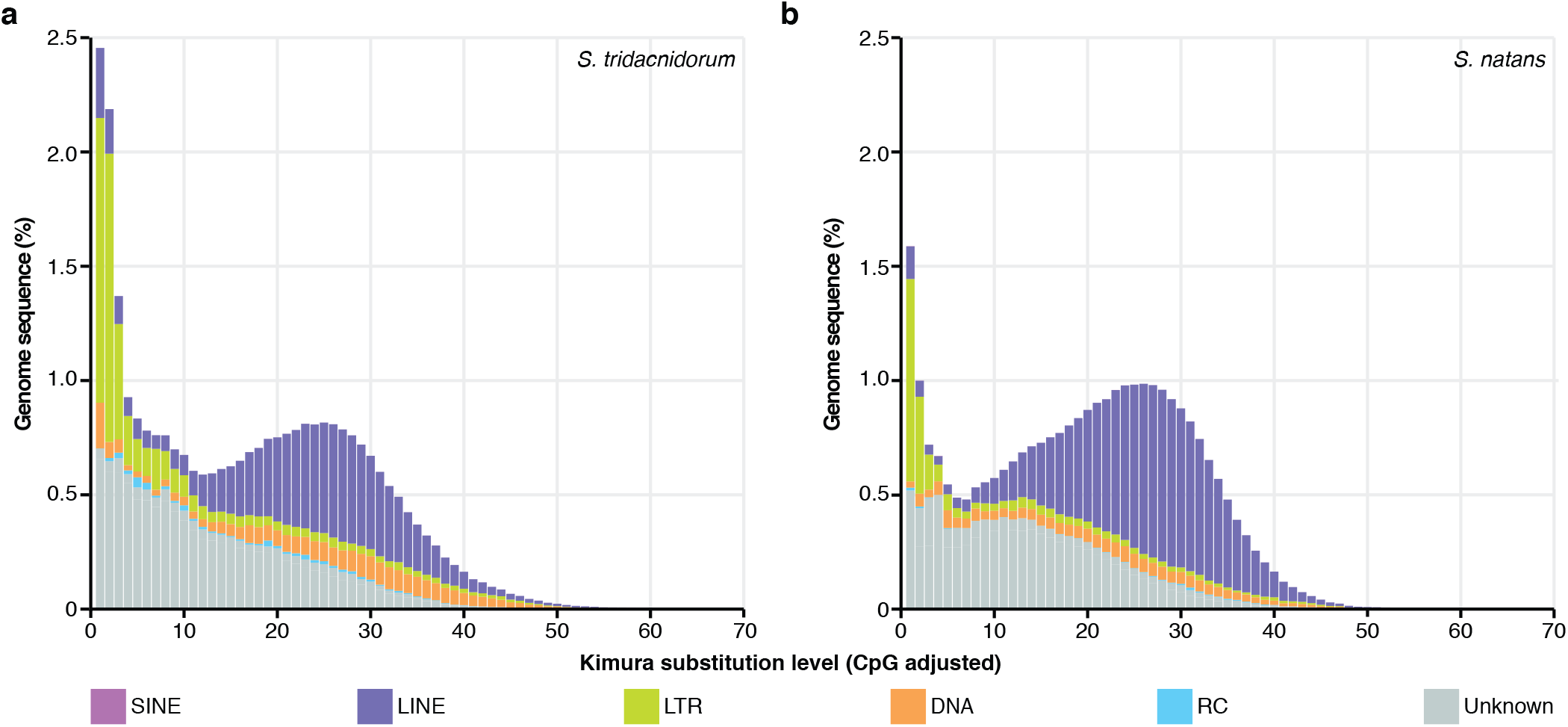
Interspersed repeat landscapes of *S. tridacnidorum* and *S. natans*. Interspersed repeat landscapes of *S. natans* **(a)** and *S. tridacnidorum* **(b)**. The colour code of the different repeat classes is shown at the bottom of the charts.

### High divergence among gene copies counteracts gene-family expansion in *S. tridacnidorum*

Duplicated genes can experience distinct fates^35,36^. These fates can result in different scenarios depending on the divergence accumulated in the sequences. First, if the function remains the same or changes slightly (*e.g.* through subfunctionalisation), the duplicated gene sequences will remain similar, resulting in gene-family expansion. We assessed the difference in gene-family sizes between *S. tridacnidorum* and *S. natans* using Fisher’s exact test (see Methods), and consider those with an adjusted *p* ≤ 0.05 as significantly different (Fig. 5). Although events contributing to the increase of gene-copy numbers appear more prevalent in *S. tridacnidorum*, gene families are not drastically larger than those in *S. natans*; only 20 families are significantly larger. Of these 20 families, one (OG0000004) putatively encodes protein kinases and glycosyltransferases that are necessary for the biosynthesis of glycoproteins, and another (OG0000013) encodes ankyrin and transport proteins (Supplementary Table 7). These functions are important for the recognition of and interaction with the host among symbiodiniacean symbionts^37–39^. In comparison, five gene families were significantly larger in *S. natans* than in *S. tridacnidorum*, of which one (OG0000003) encodes for a sodium-transporter and another (OG0000034) for a transmembrane protein. Many genes in the expanded families encode for retrotransposition functions in both genomes, lending support to the contributing role of retrotransposons in gene-family expansion in Symbiodiniaceae^31^. Although the functions of many other genes in these families could not be determined due to the lack of known similar sequences, they might be relevant for adaptation to specific ecological niches as previously proposed for dinoflagellates^40^.

**Fig. 5.**
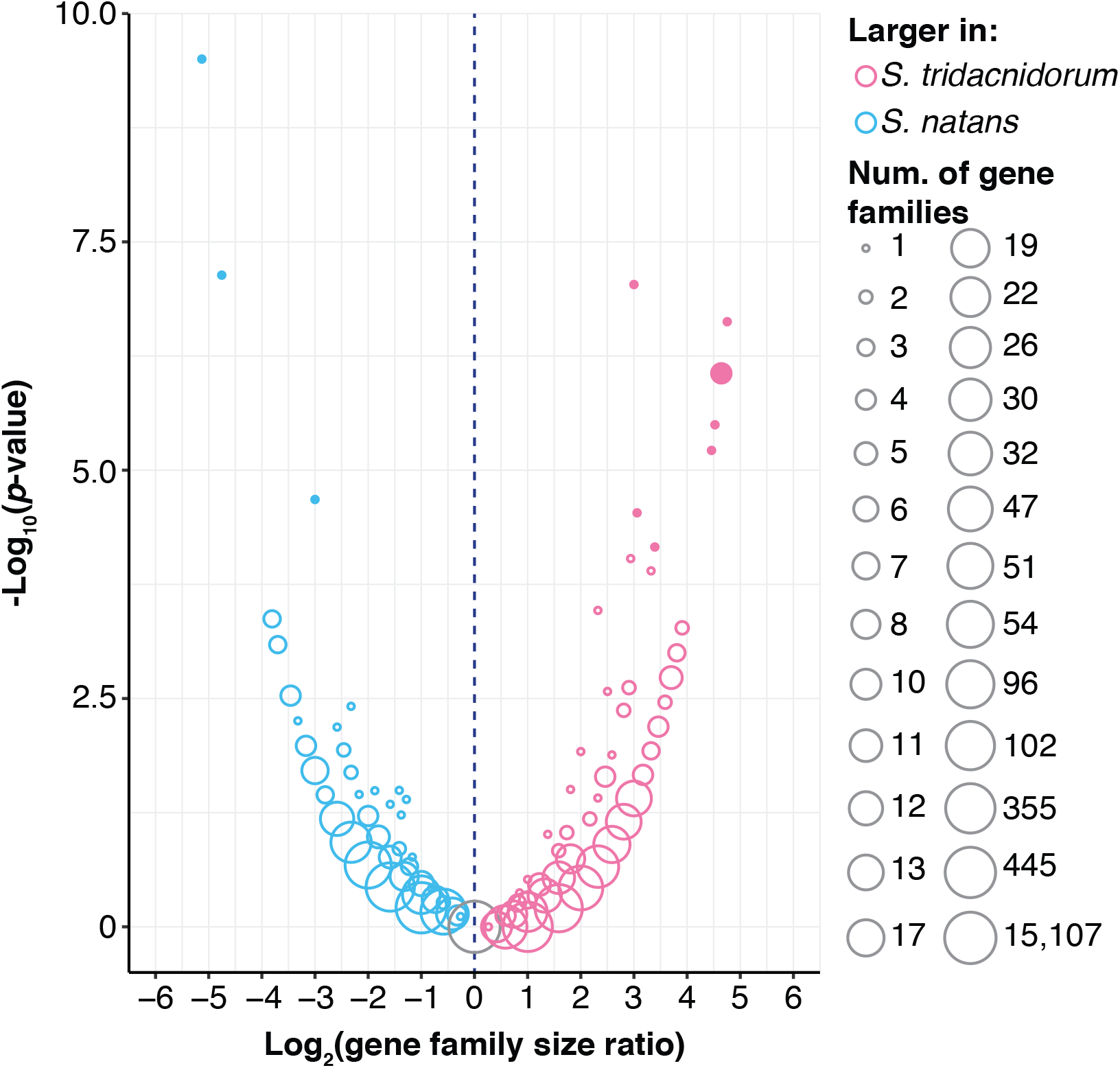
Relative gene-family sizes in *S. tridacnidorum* and *S. natans*. Volcano plot comparing gene-family sizes against Fisher’s exact test significance (*p*-value). The colour of the circles indicates the species in which those gene families are larger according to the top-right legend. The number of gene families with the same ratio and significance is represented with the circle size following the bottom-right legend. Filled circles represent size differences that are considered statistically significant (adjusted *p* ≤ 0.05).

Second, if novel beneficial functions of the gene copies emerge (*i.e.* neofunctionalisation), the sequence divergence between gene copies may become too large to be recognised as the same family. This scenario could, at least partially, explain the higher number of single-copy genes exclusive to *S. tridacnidorum* (25,649) than those exclusive to *S. natans* (16,137). Whereas 13,320 (82.54%) of the 16,137 single-copy genes of *S. natans* are supported by transcriptome evidence, only 13,189 (51.42%) of those 25,649 in *S. tridacnidorum* are. It remains unclear if these latter represent functional genes. Moreover, the annotated functions of these single-copy genes exclusive to each genome are similar in both species (Supplementary Table 8), suggesting the presence of highly diverged homologs.

Finally, duplicated genes can undergo loss of function (*i.e.* nonfunctionalisation or pseudogenisation). Pseudogene screening in both genomes (see Methods) identified 183,516 putative pseudogenes in *S. tridacnidorum* and 48,427 in *S. natans*. The nearly four-fold difference in the number of pseudogenes between the two genomes further supports the notion that more-frequent duplication events occur in *S. tridacnidorum*, and may explain the lower proportion of genes with transcript support in this species (**Error! Reference source not found.**).

Our results suggest that the high sequence divergence of duplicated genes, potentially due to the accumulation of mutations as a consequence of pseudogenisation, perhaps together with neofunctionalisation, may hinder gene family expansion in the genome of *S. tridacnidorum*.

### Gene functions of *S. tridacnidorum* and *S. natans* are relevant to their lifestyle

According to our analysis of enriched gene functions in *S. tridacnidorum* relative to *S. natans* based on annotated GO terms, methylation and the biosynthesis of histidine and peptidoglycan were among the most significant (Supplementary Table 9). The enrichment of methylation is not surprising because retrotransposons of Symbiodiniaceae are known to have acquired methyltransferase domains, likely contributing to the hypermethylated nuclear genomes of these dinoflagellates^41^. The link between the extent of methylation in symbiodiniacean genomes and its representation among predicted genes can be further assessed using methylation sequencing.

Although some corals can synthesise histidine *de novo*, metazoans generally lack this capacity^42^. The enrichment of histidine biosynthesis in *S. tridacnidorum* may be a result of host-symbiont coevolution or, alternatively, may explain why this species is a preferred symbiont over others (*e.g. S. natans*). Biosynthesis of peptidoglycans is also important for symbiosis, because these molecules, on the cell surface of Symbiodiniaceae, interact with host lectins as part of the symbiont recognition process^30,39^.

On the other hand, *S. natans* displays a wider range of enriched functions related to cellular processes (Supplementary Table 9), as expected for free-living Symbiodiniaceae^12^. One of the most significantly overrepresented gene functions is the transmembrane transport of sodium. Whereas this function is likely related to pH (osmotic) homeostasis with the extracellular environment, the occurrence of a sodium:phosphate symporter (PF02690) in tandem, exclusive to *S. natans*, and the abundance of a sodium:chloride symporter (PF00209) among the RT-genes (Supplementary Table 5) suggest that *S. natans* makes use of the Na^+^ differential gradient (caused by the higher Na^+^ concentration in seawater) for nutrient uptake in a similar fashion to the assimilation of inorganic phosphate by the malaria parasite (*Plasmodium falciparum*) in the Na^+^-rich cytosol of the host’s erythrocytes^43^.

### Are features underpinning genome divergence in Symbiodiniaceae ancestral or derived?

To assess whether the genome features found in *S. tridacnidorum* were ancestral or derived relative to *S. natans*, we compared the genome sequences from both species with those from the outgroup *Polarella glacialis* CCMP1383^22^, a psychrophilic free-living species closely related to Symbiodiniaceae (also in Order Suessiales).

A greater genome sequence proportion of *S. natans* (3.38%) than that of *S. tridacnidorum* (0.85%) aligned to the *P. glacialis* genome assembly. Interestingly, the aligned regions in both cases implicate only ~5 Mbp (~0.18%) of the *P. glacialis* genome sequence. This observation is likely due to duplicated genome regions of *S. natans* that have remained highly conserved. Similarly, the average percent identity of the best-matching sequences between any of the two *Symbiodinium* genomes against *P. glacialis* is very similar (*i.e.* 92.13% and 92.56% for *S. tridacnidorum* and *S. natans*, respectively). Nonetheless, regions occupied by duplicated genes are recovered in larger proportions in *Symbiodinium* than in *P. glacialis* (Fig. 6). On the other hand, LTR retrotransposons are evidently more prominent in *P. glacialis*. However, these LTRs are more diverged (Kimura distances 3-8)^22^ than those in the two *Symbiodinium* (Kimura distances < 5; Fig. 4), indicating an independent, more-ancient burst of these elements in *P. glacialis*.

**Fig. 6.**
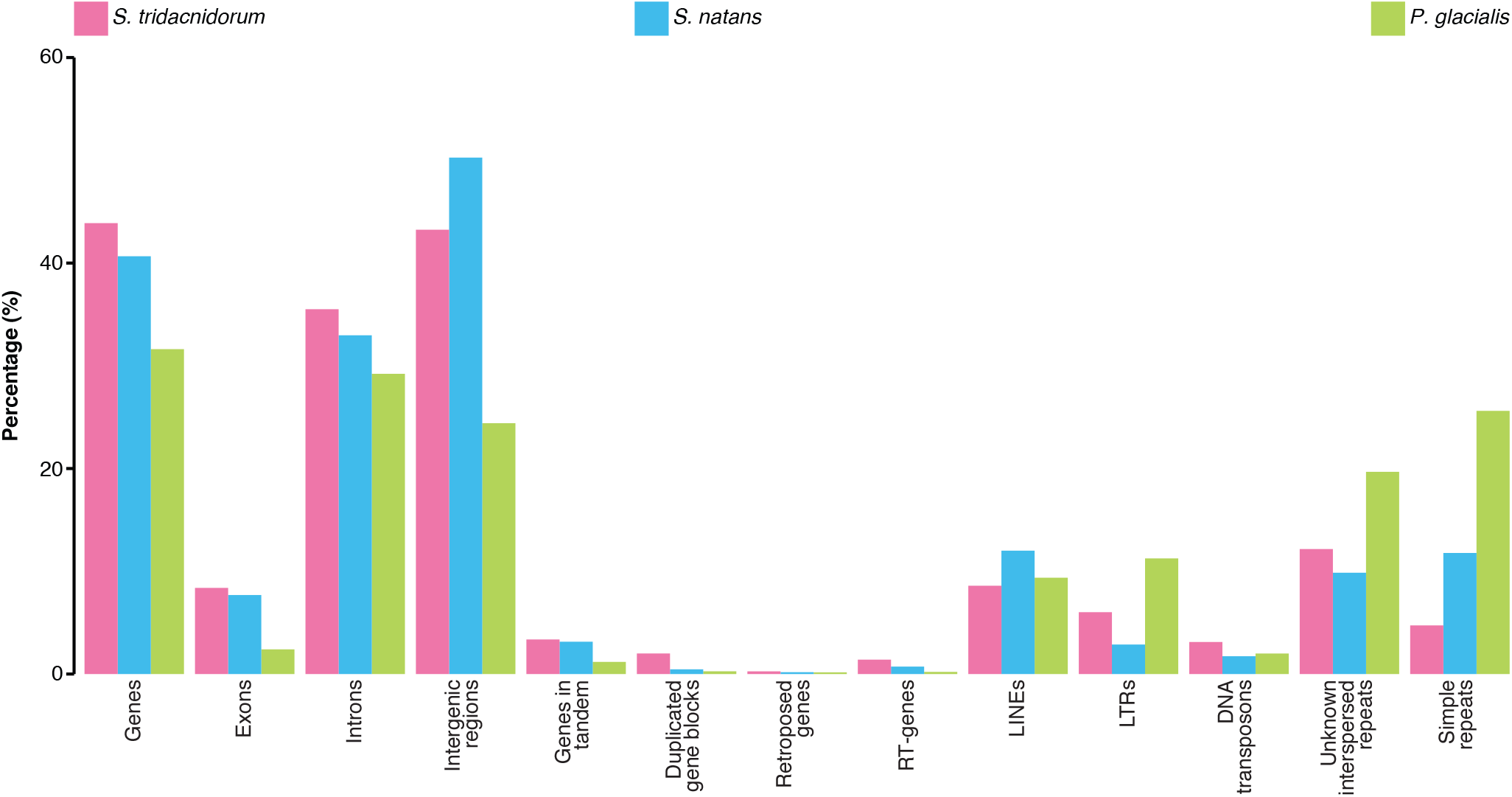
Genome proportion of distinct elements in genomes of *S. tridacnidorum*, *S. natans* and *P. glacialis*. Proportion (in percentage of the sequence length) covered by different types of genome features in the hybrid assemblies of *S. tridacnidorum*, *S. natans* and *P. glacialis*.

## Discussion

We report for the first time, based on whole-genome sequence data, evidence of structural rearrangements and TEs contributing to the extensive genomic divergence between the symbiotic *S. tridacnidorum* and the free-living *S. natans*, including the discrepancy in genome sizes. In comparison, structural rearrangements and TE activity are less prominent in the genomes of *S. natans* and the outgroup species *P. glacialis*.

Structural rearrangements, abundance of pseudogenes, and enhanced activity of TEs are common in facultative and recent intracellular symbionts and parasites^19,20^, and are expected in symbiotic Symbiodiniaceae^12^. Our results support this hypothesis. In this regard, our results agree with the notion that the symbiotic lifestyle is a derived trait in *Symbiodinium*, and that the free-living lifestyle is likely ancestral. Under this assumption, the genome proportion spanned by TEs and duplicated genes in *S. natans* is expected to be similar (if not smaller) than that in the outgroup *P. glacialis*. However, we found the proportion of duplicated genes to be larger in *S. natans* (Fig. 6), prompting two possible explanations. First, the pervasive simple repeats in the *P. glacialis* genome^22^, independently expanded along this lineage or possibly an ancestral trait in Suessiales, drastically diminishes the proportion of genic regions in the genome. Second, the free-living lifestyle of *S. natans* may be a derived trait in *Symbiodinium*, having passed through a symbiotic phase earlier in its evolutionary history. However, the robust placement of *S. natans* in the basal position alongside *Symbiodinium pilosum* (another free-living species) in the *Symbiodinium* phylogeny^2^ contradicts this less-parsimonious explanation. Additional high-quality genome data from free-living and symbiotic taxa are thus required to gain a clearer understanding of the evolutionary transition(s) between free-living and symbiotic lifestyles in Symbiodiniaceae.

## Methods

### *Symbiodinium* cultures

Single-cell monoclonal cultures of two *Symbiodinium* (formerly Clade A) species were obtained from the Bigelow National Center for Marine Algae and Microbiota. *Symbiodinium natans* (strain CCMP2548) was originally collected from open ocean water in Hawaii, USA. *Symbiodinium tridacnidorum* (Clade A3, strain CCMP2592) was originally recovered from a stony coral (*Heliofungia actiniformis*) on the Great Barrier Reef, Australia. The cultures were maintained in multiple 100-mL batches (in 250-mL Erlenmeyer flasks) in f/2 (without silica) medium (0.2 mm filter-sterilized) under a 14:10 h light-dark cycle (90 µE/m^2^/s) at 25 ºC. The medium was supplemented with antibiotics (ampicillin [10 mg/mL], kanamycin [5 mg/mL] and streptomycin [10 mg/mL]) to reduce bacterial growth.

### Nucleic acid extraction

Genomic DNA was extracted following the 2×CTAB protocol with modifications. *Symbiodinium* cells were first harvested during exponential growth phase (before reaching 10^6^ cells/mL) by centrifugation (3000 *g*, 15 min, room temperature (RT)). Upon removal of residual medium, the cells were snap-frozen in liquid nitrogen prior to DNA extraction, or stored at −80 °C. For DNA extraction, the cells were suspended in a lysis extraction buffer (400 µL; 100 mM Tris-Cl pH 8, 20 mM EDTA pH 8, 1.4 M NaCl), before silica beads were added. In a freeze-thaw cycle, the mixture was vortexed at high speed (2 min), and immediately snap-frozen in liquid nitrogen; the cycle was repeated 5 times. The final volume of the mixture was made up to 2% w/v CTAB (from 10% w/v CTAB stock; kept at 37 °C). The mixture was treated with RNAse A (Invitrogen; final concentration 20 µg/mL) at 37 °C (30 min), and Proteinase K (final concentration 120 µg/mL) at 65 °C (2 h). The lysate was then subjected to standard extractions using equal volumes of phenol:chloroform:isoamyl alcohol (25:24:1 v/v; centrifugation at 14,000 *g*, 5 min, RT), and chloroform:isoamyl alcohol (24:1 v/w; centrifugation at 14,000 *g*, 5 min, RT). DNA was precipitated using pre-chilled isopropanol (gentle inversions of the tube, centrifugation at 18,000 *g*, 15 min, 4 °C). The resulting pellet was washed with pre-chilled ethanol (70% v/v), before stored in Tris-HCl (100 mM, pH 8) buffer. DNA concentration was determined with NanoDrop (Thermo Scientific), and DNA with A_230:260:280_ ≈ 1.0:2.0:1.0 was considered appropriate for sequencing. Total RNA was isolated using the RNeasy Plant Mini Kit (Qiagen) following directions of the manufacturer. RNA quality and concentration were determined with am Agilent 2100 BioAnalyzer.

### Genome sequence data generation and *de novo* assembly

In total, we generated 1021.63 Gbp (6.77 billion reads) of genome sequence data for *S. natans* and 259.57 Gbp (1.48 billion reads) for *S. tridacnidorum* (Supplementary Table 1). Short-read sequence data (2 × 150 bp reads) were generated using multiple paired-end (for both species) and mate-pair (for *S. natans* only) libraries on the Illumina HiSeq 2500 and 4000 platforms at the Australian Genome Research Facility (Melbourne) and the Translational Research Institute Australia (Brisbane). One of the paired-end libraries for *S. natans* (of insert length 250 bp) was designed such that the read-pairs of 2 × 150 bp would overlap. Genome size and sequence read coverage were estimated based on *k*-mer frequency analysis (Supplementary Table 2) as counted with Jellyfish v2.2.6, using only pared-end data.

Quality assessment of the raw paired-end data was done with FastQC v0.11.5, and subsequent processing with Trimmomatic v0.36^44^. To ensure high-quality read data for downstream analyses, the paired-end mode of Trimmomatic was run with the settings: ILLUMINACLIP:[AdapterFile]:2:30:10 LEADING:30 TRAILING:30 SLIDINGWINDOW:4:25 MINLEN:100 AVGQUAL:30; CROP and HEADCROP were run (prior to LEADING and TRAILING) when required to remove read ends with nucleotide biases. Overlapping read pairs from the library with insert size of 250 bp were merged with FLASh v1.2.11^45^. Library adapters from the mate-pair data were removed with NxTrim v0.41^46^. A preliminary *de novo* genome assembly per species was done for genome-guided transcriptome assembly (see below) with CLC Genomics Workbench v7.5.1 (<u>qiagenbioinformatics.com</u>) using default parameters and the merged pairs (for *S. natans*), the unmerged read pairs and the trim-surviving unpaired reads. The preliminary assembly of *S. natans* was further scaffolded with SSPACE v3.0^47^ and the mate-pair filtered data.

Additionally, long-read sequence data were generated on a PacBio Sequel system at the Ramaciotti Centre for Genomics (Sydney). These data and the paired-end libraries (adding up to a coverage of 152-fold for *S. natans* and 200-fold for *S. tridacnidorum*) were used for hybrid *de novo* genome assembly (Supplementary Table 1) with MaSuRCA 3.3.0^48^, following the procedure described in the manual. Except for the PacBio sub-reads, filtered to a minimum length of 5 kbp, all sequence data were input without being pre-processed, as recommended by the developer. The genome assemblies were further scaffolded with transcriptome data generated in this study (see below) using L_RNA_scaffolder^49^.

### Removal of putative microbial contaminants

To identify putative sequences from bacteria, archaea and viruses in the genome scaffolds we followed the approach of Liu *et al.*^3^. In brief, we first searched the scaffolds (BLASTn) against a database of bacterial, archaeal and viral genomes from RefSeq (release 88); hits with *E* ≤ 10^−20^ and alignment bit score ≥1000 were considered as significant. We then calculated the proportion of bases in each scaffold covered by significant hits. Next, we assessed the added length of implicated genome scaffolds across different thresholds of these proportions, and the corresponding gene models in these scaffolds as predicted from available transcripts using PASA v2.3.3^50^ (see below), with a modified script available at <u>github.com/chancx/dinoflag-alt-splice</u>) that recognises an additional donor splice site (GA), and TransDecoder v5.2.0^50^. This preliminary gene prediction was done on the repeat-masked genome using clean transcripts, as described below. The most-stringent sequence coverage (≥5%) was selected as the threshold for all samples, *i.e.* any scaffold with significant bacterial, archaeal or viral hits covering ≥5% of its length was considered as contaminant and removed from the assembly (Supplementary Figure 5).

### RNA sequence data generation and transcriptome assembly

We generated transcriptome sequence data for both *S. tridacnidorum* and *S. natans* (Supplementary Table 10). Short-read sequence data (2 × 150 bp reads) were generated using paired-end libraries on the Illumina NovaSeq 6000 platform at the Australian Genome Research Facility (Melbourne). Quality assessment of the raw paired-end data was done with FastQC v0.11.4, and subsequent processing with Trimmomatic v0.35^44^. To ensure high-quality read data for downstream analyses, the paired-end mode of Trimmomatic was run with the settings: HEADCROP:10 ILLUMINACLIP:[AdapterFile]:2:30:10 CROP:125 SLIDINGWINDOW:4:13 MINLEN:50. The surviving read pairs were further trimmed with QUADTrim v2.0.2 (<u>bitbucket.org/arobinson/quadtrim</u>) with the flags *-m 2* and *-g* to remove homopolymeric guanine repeats at the end of the reads (a systematic error of Illumina NovaSeq 6000).

Transcriptome assembly was done with Trinity v2.1.1^51^ in two modes: *de novo* and genome-guided. *De novo* transcriptome assembly was done using default parameters and the trimmed read pairs. For genome-guided assembly, high-quality read pairs were aligned to the preliminary *de novo* genome assembly using Bowtie v2.2.7^52^. Transcriptomes were then assembled with Trinity in the genome-guided mode using the alignment information, and setting the maximum intron size to 100,000 bp. Both *de novo* and genome-guided transcriptome assemblies from each sample were used for scaffolding (see above) and gene prediction (see below).

### Full-length transcript evidence for gene prediction

Full-length transcripts for *S. tridacnidorum* and *S. natans* were generated using the PacBio IsoSeq technology. All sequencing was conducted using the PacBio Sequel platform at the Institute for Molecular Bioscience (IMB) Sequencing Facility, The University of Queensland (Brisbane, Australia; Supplementary Table 10). Full-length cDNA was first synthesised and amplified using the TeloPrime Full-Length cDNA Amplification Kit (Lexogen) and TeloPrime PCR Add-on Kit (Lexogen) following the protocols provided in the product manuals. One synthesis reaction was performed for each sample using 821 ng from *S. tridacnidorum* and 1.09 µg from *S. natans* of total RNA as starting material. Next, 25 (*S. tridacnidorum*) and 23 (*S. natans*) PCR cycles were carried out for cDNA amplification. PCR products were divided into two fractions, which were purified using 0.5× (for *S. tridacnidorum*) and 1× (for *S. natans*) AMPure PB beads (Pacific Biosciences), and then pooled with equimolar quantities. The recovered 699 ng (*S. tridacnidorum*) and 761 ng (*S. natans*) of cDNA were used for sequencing library preparation with the SMRTbell Template Prep Kit 1.0 (Pacific Biosciences). The cDNA from these libraries were sequenced in two SMRT cells.

To generate the dinoflagellate spliced-leader (DinoSL) specific transcript library, 12 PCR cycles were carried out for both samples using the conserved DinoSL fragment (5′-CCGTAGCCATTTTGGCTCAAG-3′) as forward primer, the TeloPrime PCR 3′-primer as reverse primer, and the fraction of full-length cDNA purified with 0.5× (for *S. tridacnidorum*) and 1× (for *S. natans*) AMPure PB beads. The above-described PCR purification and sequencing library preparation methods were used for the DinoSL transcript libraries; cDNA from these libraries was sequenced in one SMRT cell per sample.

Due to the abundance of undesired 5′-5′ and 3′-3′ pairs, and to recover as much transcript evidence as possible for gene prediction, we followed two approaches (Supplementary Figure 6). First, the IsoSeq 3.1 workflow (<u>github.com/PacificBiosciences/IsoSeq3/blob/master/README_v3.1.md</u>) was followed. Briefly, circular consensus sequences (CCS) were generated from the subreads of each SMRT cell with ccs v3.1.0 without polishing, and setting the minimum number of subreads to generate CCS (*--minPasses*) to 1. Removal of primers was done with lima v1.8.0 in the IsoSeq mode, with a subsequent refinement step using isoseq v3.1.0. At this stage, the refined full-length transcripts of all SMRT cells (excluding those from the DinoSL library) were combined to be then clustered by similarity and polished with isoseq v3.1.0. High- and low-quality transcripts resulting from this approach were further used for gene prediction (see below).

For the second approach, we repeated the IsoSeq workflow with some modifications. We polished the subreads with the Arrow algorithm and used at least three subreads per CCS with ccs v3.1.0 to generate high-accuracy CCS. Primer removal and refinement were done as explained above. The subsequent clustering and polishing steps were skipped. The resulting polished CCS and full-length transcripts were also used for gene prediction. IsoSeq data from the DinoSL library were processed separately following the same two approaches.

### Genome annotation and gene prediction

We adopted the same comprehensive *ab initio* gene prediction approach reported in Chen *et al.*^53^, using available genes and transcriptomes of Symbiodiniaceae as guiding evidence. A *de novo* repeat library was first derived for the genome assembly using RepeatModeler v1.0.11 (<u>repeatmasker.org/RepeatModeler</u>). All repeats (including known repeats in RepeatMasker database release 20180625) were masked using RepeatMasker v4.0.7 (<u>repeatmasker.org</u>).

As direct transcript evidence, we used the *de novo* and genome-guided transcriptome assemblies from Illumina short-read sequence data, as well as the PacBio IsoSeq full-length transcript data (see above). We concatenated all the transcript datasets per sample and “cleaned” them with SeqClean (<u>sourceforge.net/projects/seqclean</u>) and the UniVec database build 10.0. We used PASA v2.3.3^50^, customised to recognise dinoflagellate alternative splice donor sites (see above), and TransDecoder v5.2.0^50^ to predict coding sequences (CDS). These CDS were searched (BLASTp, *E* ≤ 10^−20^) against a protein database that consists of RefSeq proteins (release 88) and a collection of available and predicted (with TransDecoder v5.2.0^50^) proteins of Symbiodiniaceae (total of 111,591,828 sequences; Supplementary Table 11). We used the *analyze_blastPlus_topHit_coverage.pl* script from Trinity v2.6.6^51^ to retrieve only those CDS having a hit with >70% coverage of the database protein sequence (*i.e.* nearly full-length) in the database for subsequent analyses.

The near full-length gene models were checked for TEs using HHblits v2.0.16 (probability = 80% and *E*-value = 10^−5^), searching against the JAMg transposon database (<u>sourceforge.net/projects/jamg/files/databases</u>), and TransposonPSI (<u>transposonpsi.sourceforge.net</u>).

Gene models containing TEs were removed from the gene set, and redundancy reduction was conducted using cd-hit v4.6^54,55^ (ID = 75%). The remaining gene models were processed using the *prepare_golden_genes_for_predictors.pl* script from the JAMg pipeline (altered to recognise GA donor splice sites; <u>jamg.sourceforge.net</u>). This script produces a set of “golden genes” that was used as training set for the *ab initio* gene-prediction tools AUGUSTUS v3.3.1^56^ (customised to recognise the non-canonical splice sites of dinoflagellates, following the changes made to that available at <u>smic.reefgenomics.org/download</u>) and SNAP v2006-07-28^57^. Independently, the soft-masked genome sequences were passed to GeneMark-ES v4.32^58^ for unsupervised training and gene prediction. UniProt-SwissProt proteins (downloaded on 27 June 2018) and the predicted proteins of Symbiodiniaceae (Supplementary Table 11) were used to produce a set of gene predictions using MAKER v2.31.10^59^ protein2genome; the custom repeat library was used by RepeatMasker as part of MAKER prediction. A primary set of predicted genes was produced using EvidenceModeler v1.1.1^60^, modified to recognise GA donor splice sites. This package combined the gene predictions from PASA, SNAP, AUGUSTUS, GeneMark-ES and MAKER protein2genome into a single set of evidence-based predictions. The weightings used for the package were: PASA 10, Maker protein 8, AUGUSTUS 6, SNAP 2 and GeneMark-ES 2. Only gene models with transcript evidence (*i.e.* predicted by PASA) or supported by at least two *ab initio* prediction programs were kept. We assessed completeness by querying the predicted protein sequences in a BLASTp similarity search (E ≤ 10^−5^, ≥50% query/target sequence cover) against the 458 core eukaryotic genes from CEGMA^21^. Transcript data support for the predicted genes was determined by BLASTn (*E* ≤ 10^−5^) similarity search, querying the transcript sequences against the predicted CDS from each genome. Genes for which the transcripts aligned to their CDS with at least 50% of sequence cover and 90% identity were considered as supported by transcript data.

### Gene-function annotation and enrichment analyses

Annotation of the predicted genes was done based on sequence similarity searches against know proteins following the same approach as Liu *et al.*^3^, in which the predicted protein sequences were used as query (BLASTp, *E* ≤ 10^−5^, minimum query or target cover of 50%) against Swiss-Prot first, and those with no Swiss-Prot hits subsequently against TrEMBL (both databases from UniProt, downloaded on 27 June 2018). The best UniProt hit with associated Gene Ontology (GO, <u>geneontology.org</u>) terms was used to annotate the query protein with those GO terms using the UniProt-GOA mapping (downloaded on 03/06/2019). Pfam domains^61^ were searched in the predicted proteins of both *Symbiodinium* species using PfamScan^62^ (*E* ≤ 0.001) and the Pfam-A database (release 30 August 2018)^61^.

Tests for enrichment of Pfam domains were done with one-tailed Fisher’s exact tests, independently for over- and under-represented features; domains with Benjamini-Hochberg^63^ adjusted p ≤ 0.05 were considered significant. Enrichment of GO terms was performed using the topGO Bioconductor package^64^ implemented in R v3.5.1, applying Fisher’s Exact test with the ‘elimination’ method to correct for the dependence structure among GO terms. GO terms with a p ≤ 0.01 were considered significant.

### Comparative genomic analyses

Whole-genome sequence alignment was carried out with nucmer v4.0.0^65^ with the hybrid genome assembly of *S. natans* as reference and that of *S. tridacnidorum* as query, and using anchor matches that are unique in the sequences from both species (*--mum*). Sequences from both *Symbiodinium* genomes were queried in the same way against the genome sequence of *P. glacialis* CCMP1383^22^. Filtered read pairs (see above, Supplementary Table 1) from both species were aligned to their corresponding and counterpart genome sequences using bwa v0.7.13^66^, and rates of mapping with different quality scores were calculated with SAMStat v1.5.1^67^.

Groups of homologous sequences from the two *Symbiodinium* genomes were inferred with Orthofinder v2.3.1^68^, and considered gene families. The significance of size differences of the gene families shared by *S. tridacnidorum* and *S. natans* was assessed with a two-tailed Fisher’s exact test correcting p-values for multiple testing with the Benjamini-Hochberg method^63^; difference in size was considered significant for gene families with adjusted *p* ≤ 0.05.

We used the predicted genes and their associated genomic positions to identify potential segmental genome duplications in both *Symbiodinium* species, as well as in *P. glacialis*. First, we used BLASTp (*E* ≤ 10^−5^) to search for similar proteins within each genome; the hit pairs were filtered to include only those where the alignment covered at least half of either the query or the matched protein sequence. Next, we ran MCScanX^69^ in intra-specific mode (*-b 1*) to identify collinear syntenic blocks of at least five genes and genes arranged in tandem within each genome separately.

Identification of genes with DinoSL and pseudogenes was done in a similar way to Song *et al.* (2017)^27^. We queried the original DinoSL sequence (DCCGUAGCCAUUUUGGCUCAAG)^28^, excluding the first ambiguous position, against the upstream regions (up to 500 bp) of all genes in a BLASTn search, keeping the default values of all alignment parameters but with word size set to 9 (*-word_size 9*). Pseudogene detection was done with tBLASTn, with the predicted protein for each genome as query against the genome sequence, with the regions covered by the predicted genes masked, as target. Matched regions with ≥75% identity were considered part of pseudogenes and surrounding matching fragments were considered as part of the same pseudogene as long as they were at a maximum distance of 1 kbp from another pseudogene fragment and in the same orientation.

## Acknowledgements

R.A.G.P. is supported by an International Postgraduate Research Scholarship and a University of Queensland Centenary Scholarship. This work is supported by two Australian Research Council grants (DP150101875 awarded to M.A.R., C.X.C. and D.B., and DP190102474 awarded to C.X.C. and D.B.), and the computational resources of the National Computational Infrastructure (NCI) National Facility systems through the NCI Merit Allocation Scheme (Project d85) awarded to C.X.C. and M.A.R.

## Author contributions

R.A.G.P., M.A.R. and C.X.C. conceived the study; R.A.G.P., T.G.S., A.R.M., D.W.B., D.B., M.A.R. and C.X.C. designed the analyses and interpreted the results; C.X.C. maintained the dinoflagellate cultures; C.X.C. and A.R.M. extracted biological materials for sequencing; Y. Cheng generated the long-read libraries for genome and full-length transcriptome sequencing; R.A.G.P. and Y. Chen conducted all computational analyses. R.A.G.P. prepared all figures and tables, and prepared the first draft of the manuscript; all authors wrote, reviewed, commented on and approved the final manuscript.

## Data availability

The assembled genomes, predicted gene models and proteins from *S. tridacnidorum* CCMP2592 and *S. natans* CCMP2548 are available at https://cloudstor.aarnet.edu.au/plus/s/095Tqepmq2VBztd.

---

**Table 1.** Statistics of *de novo* genome assemblies of *S. tridacnidorum* CCMP2592 and *S. natans*

| Metric | *S. tridacnidorum* | *S. natans* |
| --- | --- | --- |
| Overall G+C (%) | 51.01 | 51.79 |
| Number of scaffolds | 6245 | 2855 |
| Assembly length (bp) | 1,103,301,044 | 761,619,964 |
| N50 scaffold length (bp) | 651,264 | 610,496 |
| Max. scaffold length (Mbp) | 4.01 | 3.40 |
| Number of contigs (bp) | 7913 | 4262 |
| N50 contig length (bp) | 356,695 | 358,021 |
| Max. contig length (Mbp) | 2.96 | 2.90 |
| Gap (%) | 0.02 | 0.02 |

**Table 2.** Statistics of predicted genes from genomes of *S. tridacnidorum* and *S. natans*.

| Statistic |  | *S. tridacnidorum* | *S. natans* |
| --- | --- | --- | --- |
| **Genes** |  |  |  |
| Number of genes |  | 45,474 | 35,270 |
| Mean gene (exons + introns) length (bp) |  | 10647.95 | 8779.96 |
| Mean CDS length (bp) |  | 2033.50 | 1660.13 |
| Gene content (total gene length/total assembly length, %) |  | 43.87 | 40.66 |
| CDS G+C (%) |  | 57.32 | 58.16 |
| Supported by transcript data (%) |  | 61.73 | 82.99 |
| **Exons** |  |  |  |
| Average number per gene |  | 16.15 | 15.66 |
| Average length (bp) |  | 125.89 | 106.00 |
| Total length (bp) |  | 92,471,373 | 58,552,877 |
| **Introns** |  |  |  |
| Number of genes with introns |  | 40,282 | 30,171 |
| Average length |  | 568.48 | 485.61 |
| Total length (bp) |  | 391,733,376 | 251,116,222 |
| G+C (%) |  | 50.20 | 51.33 |
| **Intron-exon boundaries** |  |  |  |
| 5'-donor splice sites (%) | GC (canonical) | 56.38 | 58.04 |
|  | GT (non-canonical) | 25.71 | 23.60 |
|  | GA (non-canonical) | 17.91 | 18.36 |
| Nucleotide after the AG 3'-acceptor splice sites (%) | G | 96.53 | 97.09 |
|  | A | 1.98 | 1.75 |
|  | T | 0.92 | 0.78 |
|  | C | 0.57 | 0.38 |
| **Intergenic regions** |  |  |  |
| Average length (bp) |  | 11,467.68 | 11,585.13 |
| G+C (%) |  | 50.20 | 51.50 |

