## Supplementary Figures for "Structural rearrangements drive extensive genome divergence between symbiotic and free-living *Symbiodinium*"

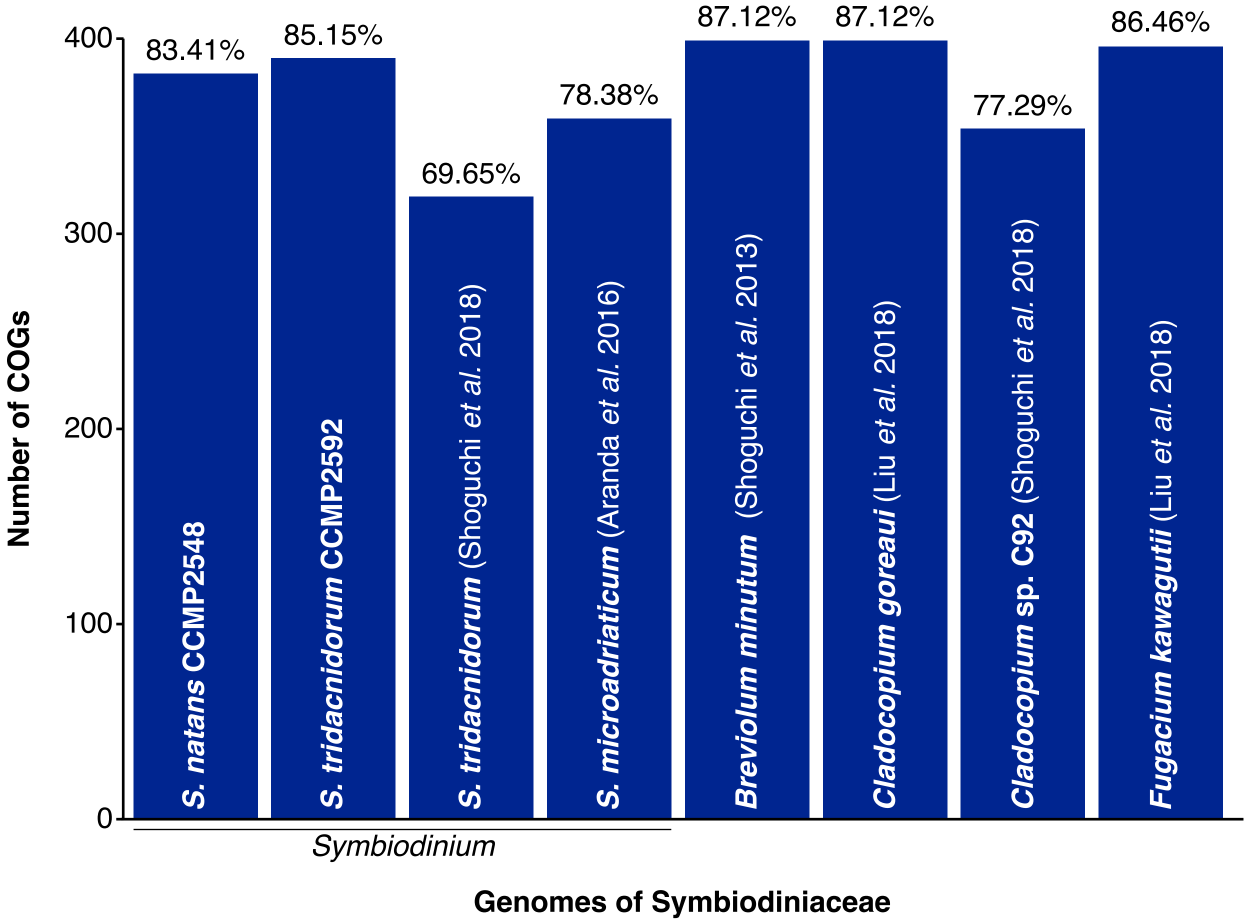


**Supplementary Figure 1** Recovery of CEGMA Clusters of Orthologous Groups (COGs) by BLASTp similarity search (see Methods) in the predicted protein sequences of *S. natans* CCMP2548 and *S. tridacnidorum* CCMP2592, as well as in the most recently predicted protein sequences[1] of other available genomes of Symbiodiniaceae: *S. tridacnidorum*[2], *Symbiodinium microadriaticum*[3], *Breviolum minutum*[4], *Cladocopium goreaui*[5], *Cladocopium* sp. C92[2] and *F. kawagutii*[5]. The fraction of COGs for each genome, out of the 458 in CEGMA, is shown on top of its corresponding bar.

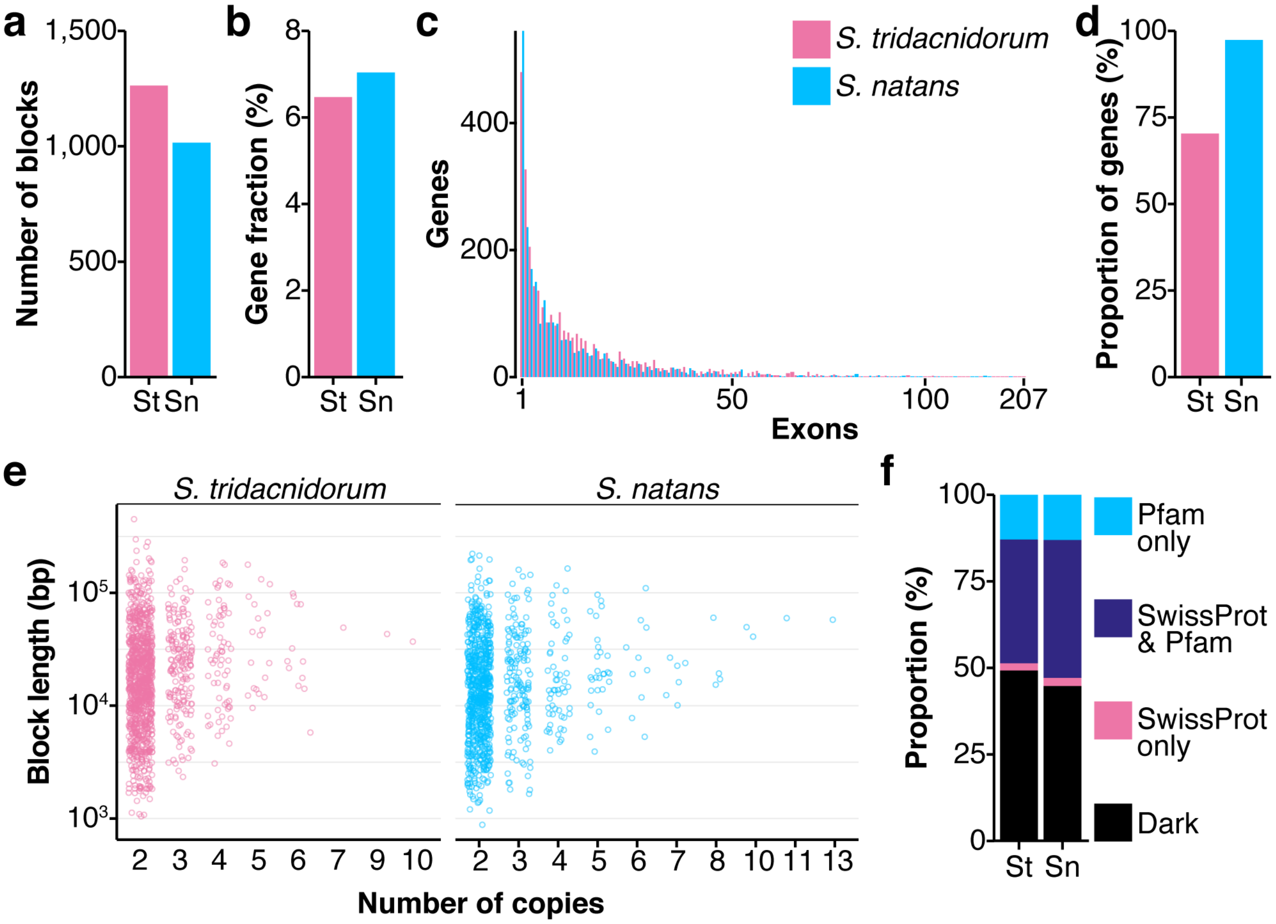


**Supplementary Figure 2** **(a)** Number of blocks of genes arranged in tandem, and **(b)** fraction of genes (out of the total number of genes per genome) implicated in those blocks, found in genomes of *S. tridacnidorum* and *S. natans*. **(c)** Breakdown of the number of genes by the number of exons they contain. **(d)** Gene fraction (out of the total number of genes in tandem) with transcript support. **(e)** Block length distribution of genes in tandem as a function of the number of gene copies implicated in the blocks. **(f)** Fraction of genes with functional annotation based either on sequence similarity of their protein products with sequences in UniProt-SwissProt or identified Pfam protein domains, or both.

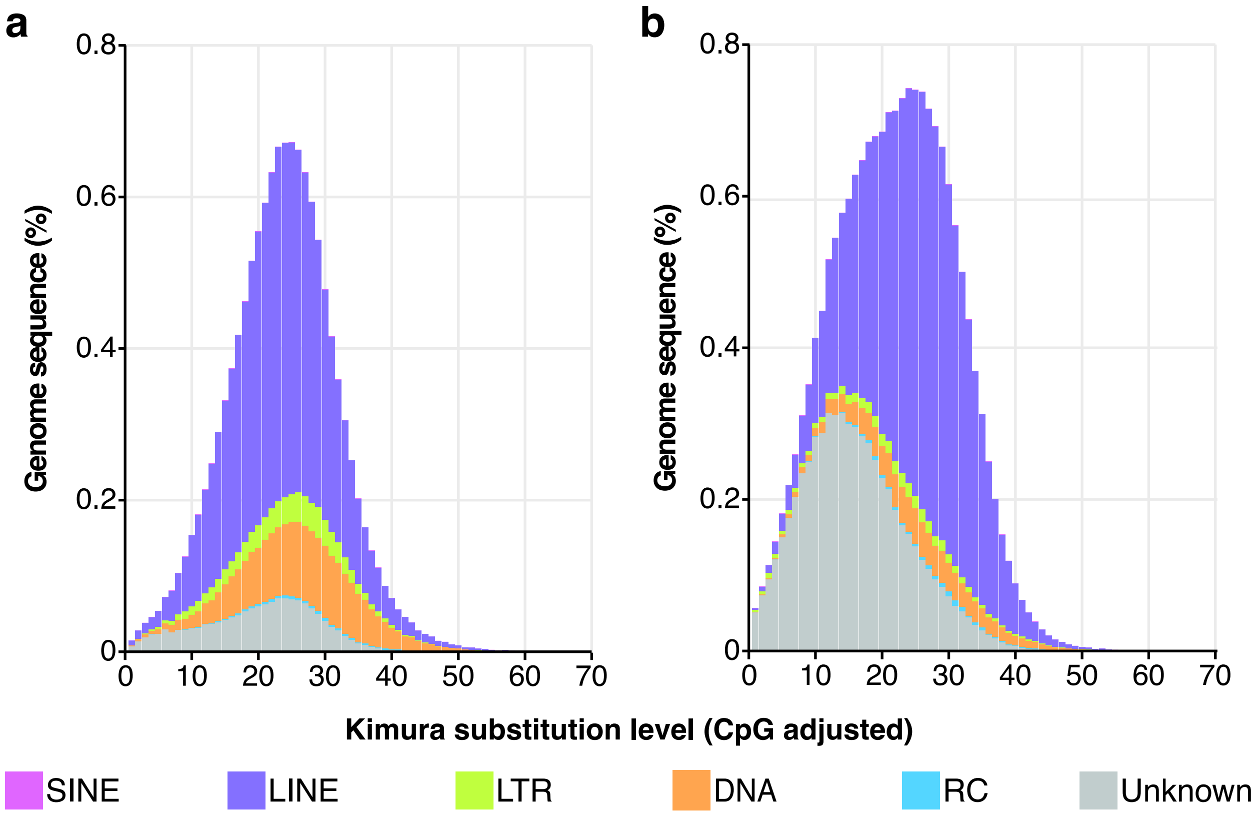


**Supplementary Figure 3** Interspersed repeat landscapes from the preliminary genome assemblies (based on short-read data only) of *S. natans* **(a)** and *S. tridacnidorum* CCMP2592 **(b)**. The colour code of the different repeat classes (including multiple-copy genes) is shown at the bottom of the two charts.

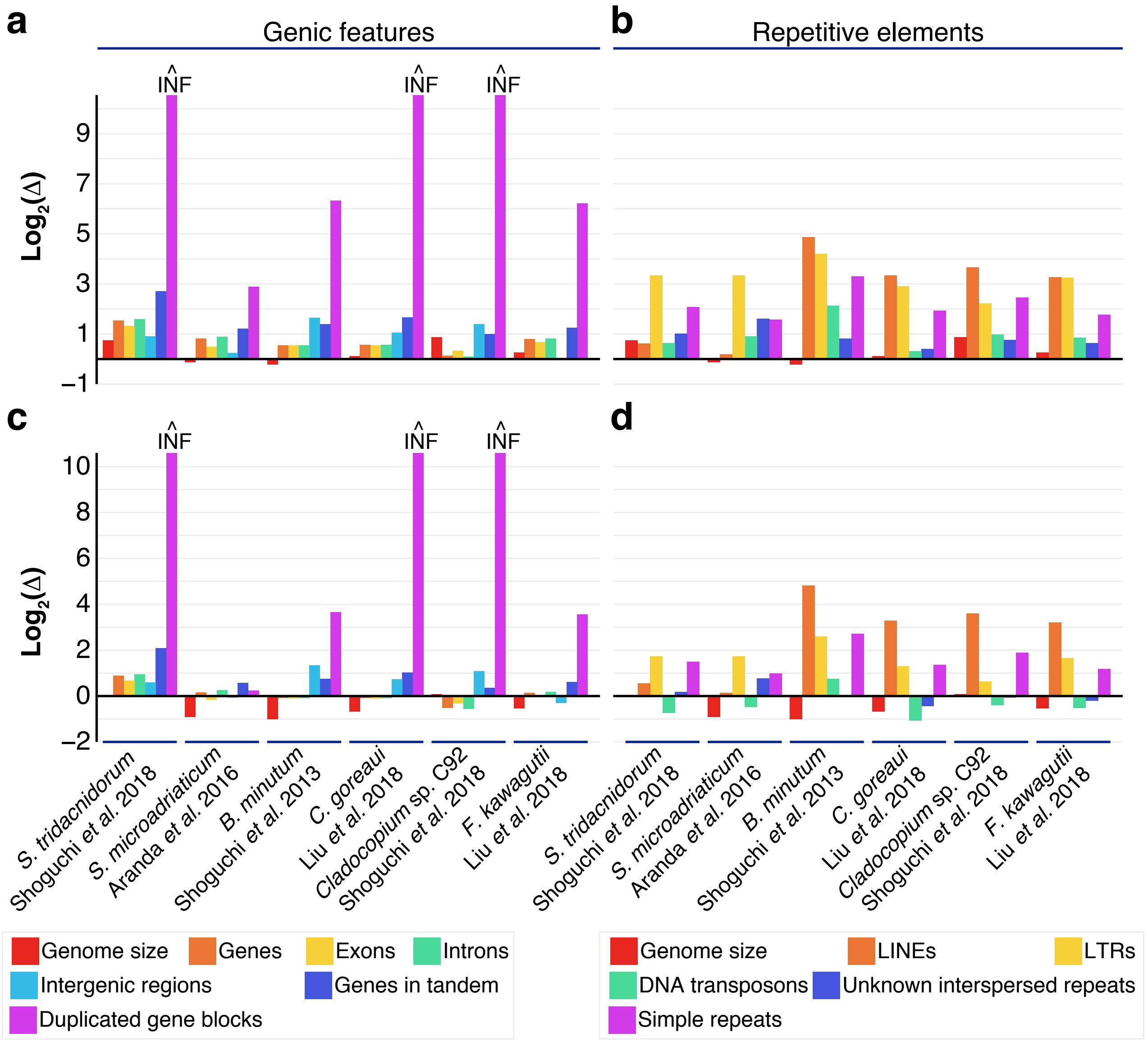


**Supplementary Figure 4** Ratio (Δ) of the sequence length covered by genic features **(a, c)** and repetitive elements **(b, d)** from other Symbiodiniaceae genome assemblies (based on short-sequence read data) relative to those from the hybrid assemblies of *S. tridacnidorum* **(a, b)** and *S. natans* **(c, d)**. The ratios between estimated genome sizes is given as reference, except for *S. tridacnidorum* and *Cladocopium* sp. C92 (Shoguchi *et al.* 2018)[2] that lack estimates, for which assembly total length was used instead.

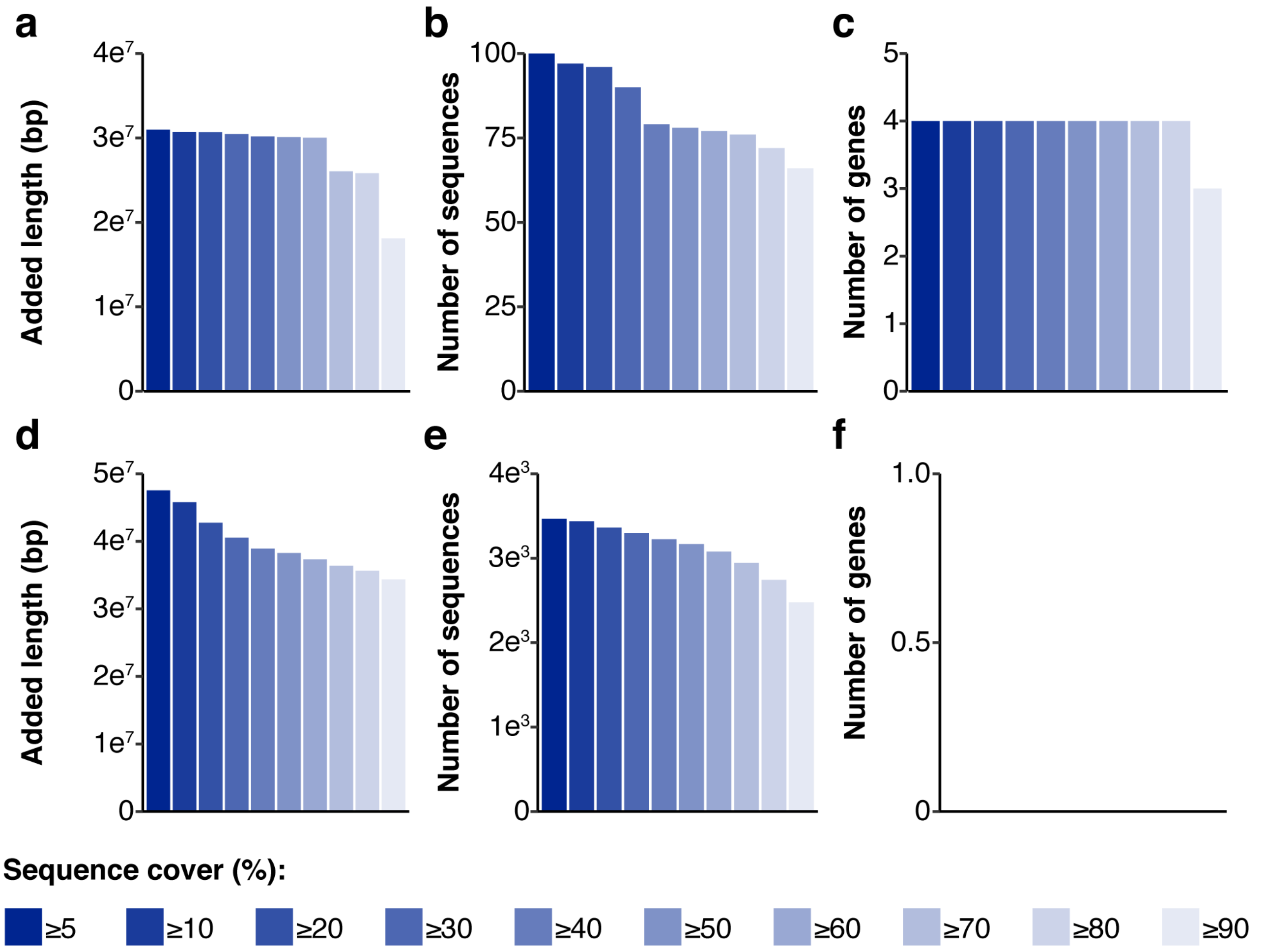


**Supplementary Figure 5** Added length **(a, d)** and count **(b, e)** of putatively contaminant sequences, and number of implicated genes **(c, f)**, in genomes assemblies from *S. tridacnidorum* **(a-c)** and *S. natans* **(d-f)**, based on different sequence cover thresholds (x-axis) of significant hits against bacterial, archaeal and viral genomes. The bars in each chart are coloured according to their corresponding query cover threshold following the colour code at the bottom.

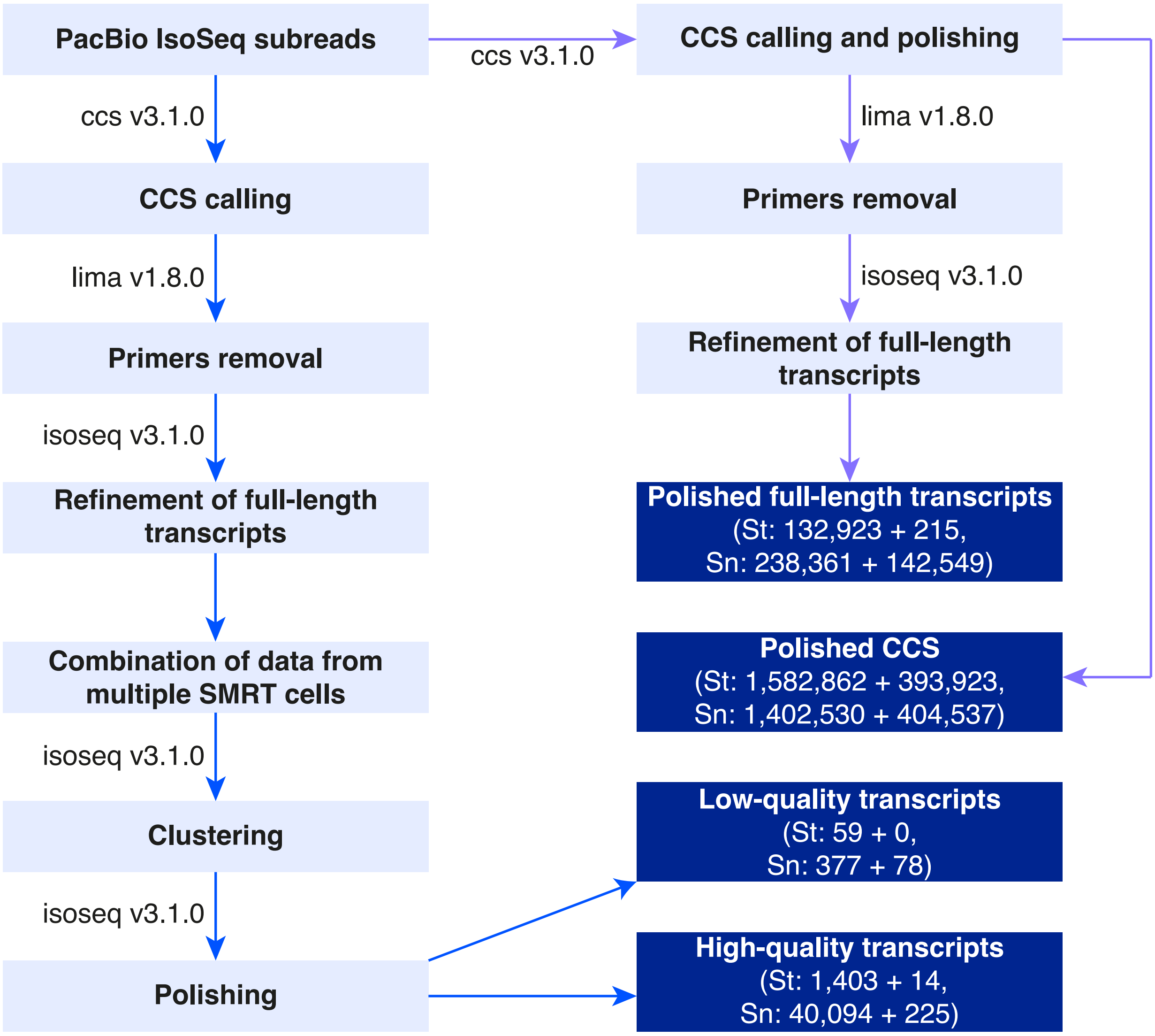


**Supplementary Figure 6** Diagram showing the detailed steps followed to process PacBio IsoSeq data to generate full-length transcript evidence for gene prediction. The traditional IsoSeq 3.1 workflow (blue arrows) was followed to obtain low- and high- quality transcripts. In an alternative approach (purple arrows), circular consensus sequences (CCS) were called and polished simultaneously. These polished CCS were further trimmed and refined into full-length transcripts skipping the clustering step. IsoSeq sequences from the DinoSL library were processed apart from the other libraries. Boxes in dark blue represent the transcript evidence subsequently used for gene prediction and the values in parentheses show the corresponding number of sequences from the standard (left) + the DinoSL (right) libraries for both *S. tridacnidorum* (St) and *S. natans* (Sn).
